# The selective autophagy receptors Optineurin and p62 are both required for innate host defense against mycobacterial infection

**DOI:** 10.1101/415463

**Authors:** Rui Zhang, Monica Varela, Wies Vallentgoed, Michiel van der Vaart, Annemarie H. Meijer

## Abstract

Mycobacterial pathogens are the causative agents of chronic infectious diseases like tuberculosis and leprosy. Autophagy has recently emerged as an innate mechanism for defense against these intracellular pathogens. *In vitro* studies have shown that mycobacteria escaping from phagosomes into the cytosol are ubiquitinated and targeted by selective autophagy receptors. However, there is currently no *in vivo* evidence for the role of selective autophagy receptors in defense against mycobacteria, and the importance of autophagy in control of mycobacterial diseases remains controversial. Here we have used *Mycobacterium marinum* (Mm), which causes a tuberculosis-like disease in zebrafish, to investigate the function of two selective autophagy receptors, Optineurin (Optn) and SQSTM1 (p62), in host defense against a mycobacterial pathogen. To visualize the autophagy response to Mm *in vivo*, *optn* and *p62* zebrafish mutant lines were generated in the background of a GFP-Lc3 autophagy reporter line. We found that loss-of-function mutation of *optn* or *p62* reduces autophagic targeting of Mm, and increases susceptibility of the zebrafish host to Mm infection. Transient knockdown studies confirmed the requirement of both selective autophagy receptors for host resistance against Mm infection. For gain-of-function analysis, we overexpressed *optn* or *p62* by mRNA injection and found this to increase the levels of GFP-Lc3 puncta in association with Mm and to reduce the Mm infection burden. Taken together, our results demonstrate that both Optineurin and p62 are required for autophagic host defense against mycobacterial infection and support that protection against tuberculosis disease may be achieved by therapeutic strategies that enhance selective autophagy.

**Author summary:** Tuberculosis is a serious infectious disease that claims over a million lives annually. Vaccination provides insufficient protection and the causative bacterial pathogen, *Mycobacterium tuberculosis*, is becoming increasingly resistant to antibiotic therapy. Therefore, there is an urgent need for novel therapeutic strategies. Besides searches for new antibiotics, considerable efforts are being made to identify drugs that improve the immune defenses of the infected host. One host defense pathway under investigation for therapeutic targeting is autophagy, a cellular housekeeping mechanism that can direct intracellular bacteria to degradation. However, evidence for the anti-mycobacterial function of autophagy is largely based on studies in cultured cells. Therefore, we set out to investigate anti-mycobacterial autophagy using zebrafish embryos, which develop hallmarks of tuberculosis following infection with *Mycobacterium marinum*. Using red-fluorescent mycobacteria and a green-fluorescent zebrafish autophagy reporter we could visualize the anti-mycobacterial autophagy response in a living host. We generated mutant and knockdown zebrafish for two selective autophagy receptors, Optineurin and p62, and found that these have reduced anti-bacterial autophagy and are more susceptible to tuberculosis. Moreover, we found that increased expression of these receptors enhances anti-bacterial autophagy and protects against tuberculosis. These results provide new evidence for the host-protective function of selective autophagy in tuberculosis.

## Introduction

Autophagy is a fundamental cellular pathway in eukaryotes that functions to maintain homeostasis by degradation of cytoplasmic contents in lysosomes (1). During autophagy, protein aggregates or defective organelles are sequestered by double-membrane structures, called isolation membranes or phagophores, which mature into autophagosomes capable of fusing with lysosomes. Autophagy was previously considered a strictly non-selective bulk degradation pathway. However, recent comprehensive studies have highlighted its selective ability. Selective autophagy depends on receptors that interact simultaneously with the cytoplasmic material and with the autophagosome marker microtubule-associated protein 1 light chain 3 (Lc3), thereby physically linking the cargo with the autophagy compartment (2, 3). Different selective autophagy pathways are classified according to their specific cargo; for example, mitophagy is the pathway that degrades mitochondria, aggrephagy targets misfolded proteins or damaged organelles, and xenophagy is directed against intracellular microorganisms. Recent studies have firmly established xenophagy as an effector arm of the innate immune system (4-6). The xenophagy pathway targets microbial invaders upon their escape from phagosomes into the cytosol, where they are coated by ubiquitin. These ubiquitinated microbes are then recognized by selective autophagy receptors of the Sequestosome (p62/SQSTM1)-like receptor (SLR) family, including p62, Optineurin, NDP52, NBRC1, and TAX1BP1 (5). In addition to targeting microbes to autophagy, SLRs also deliver ubiquitinated proteins to the same compartments. It has been shown that the processing of these proteins into neo-antimicrobial peptides is important for elimination of the pathogen *Mycobacterium tuberculosis* in macrophages (7).

*M. tuberculosis* (Mtb) is the causative agent of chronic and acute tuberculosis (Tb) infections that remain a formidable threat to global health, since approximately one-third of the human population carry latent infections and 9 million new cases of active disease manifest annually. Current therapeutic interventions are complicated by increased incidence of multi-antibiotic resistance of Mtb and co-infections with Human Immunodeficiency Virus (HIV). Despite decades of extensive research efforts, the mechanisms of how Mtb subverts the host’s innate immune defenses are incompletely understood, which poses a bottleneck for developing novel therapeutic strategies (8). Because of the discovery of autophagy as an innate host defense mechanism, the potential of autophagy-inducing drugs as adjunctive therapy for Tb is now being explored (9).

Many studies have shown that induction of autophagy in macrophages by starvation, interferon-γ (IFN-γ) treatment, or by autophagy-inducing drugs, promotes maturation of mycobacteria-containing phagosomes and increases lysosome-mediated bacterial killing (7, 10-12). Furthermore, it has been shown that the ubiquitin ligase Parkin and the ubiquitin-recognizing SLRs p62 and NDP52 are activated by the escape of Mtb from phagosomes into the cytosol (13, 14). Subsequently, the ubiquitin-mediated xenophagy pathway targets Mtb to autophagosomes (13, 14). Parkin-deficient mice are extremely vulnerable to Mtb infection (14). However, a recent study has questioned the function of autophagy in the host immune response against Mtb, since mutations in several autophagy proteins, with the exception of ATG5, did not affect the susceptibility of mice to acute Mtb infection (15). The susceptibility of ATG5-deficient mice in this study was attributed to the ability of ATG5 to prevent a neutrophil-mediated immunopathological response rather than to direct autophagic elimination of Mtb. In the same study, loss of p62 did not affect the susceptibility of mice to Tb, despite that p62 has previously been shown to be required for autophagic control of Mtb in macrophages (7, 15). These different reports suggest that Mtb employs virulence mechanisms to suppress autophagic defense mechanisms and that the host requires autophagy induction as a countermeasure (12). Taken together, the role that autophagy plays in Tb is complex and further studies are required to determine if pharmacological intervention in this process is useful for a more effective control of this disease.

In this study, we utilized zebrafish embryos and larvae to investigate the role of selective autophagy during the early stages of mycobacterial infection, prior to the activation of adaptive immunity. Zebrafish is a well-established animal model for Tb that has generated important insights into host and bacterial factors determining the disease outcome (16, 17). Infection of zebrafish embryos with *Mycobacterium marinum* (Mm), a pathogen that shares the majority of its virulence factors with Mtb, results in the formation of granulomatous aggregates of infected macrophages, considered as a pathological hallmark of Tb (17-19). Using a combination of confocal imaging in GFP-Lc3 transgenic zebrafish and transmission electron microscopy, we have previously shown that the autophagy machinery is activated during the early stages of granuloma formation in this model (20, 21). Furthermore, we found that the DNA-damage regulated autophagy modulator Dram1 protects the zebrafish host against Mm infection by a p62-dependent mechanism (21). However, the role of p62 and other SLRs in host defense against Mm remains to be further elucidated.

p62 is known to function cooperatively with Optineurin in xenophagy of *Salmonella enterica* (22-24). Both these SLRs are phosphorylated by Tank-binding kinase 1 (TBK1) and bind to different microdomains of ubiquitinated bacteria as well as interacting with Lc3 (23, 25). While several studies have implicated p62 in autophagic defense against Mtb, Optineurin has thus far not been linked to control of mycobacterial infection (7, 13, 24-26). We found gene expression of *p62* and *optn* to be coordinately upregulated during granuloma formation in zebrafish larvae (27), and set out to study the function of these SLRs by CRISPR/Cas9-mediated mutagenesis. We found that either p62 or Optineurin deficiency increased the susceptibility of zebrafish embryos to Mm infection, while overexpression of *p62* or *optn* mRNAs enhanced Lc3 association with Mm and had a host-protective effect. These results provide new *in vivo* evidence for the role of selective autophagy as an innate host defense mechanism against mycobacterial infection.

## Results

### *Mycobacterium marinum* bacteria are ubiquitinated during infection of zebrafish

Phagosomal permeabilization and cytosolic escape of Mtb is known to induce the STING-dependent DNA-sensing pathway, resulting in ubiquitination and targeting of bacteria to autophagy (13). We have previously shown that this pathway is also functional in zebrafish larvae infected with Mm and that a failure to induce autophagy reduces host resistance (21). However, it had not been formally demonstrated that Mm bacteria are ubiquitinated in this model. To examine whether ubiquitin interacts with Mm and Lc3 during infection of zebrafish, we infected embryos at 28 hours post fertilization (hpf) and performed immunostaining for ubiquitin at 1, 2, and 3 days post-infection (dpi), time points at which the early stages of tuberculous granuloma formation can be observed (Fig1 A). This process of granuloma formation is known to be induced by infected macrophages, which attract new macrophages that subsequently also become infected (28). Developing granulomas also attract neutrophils and usually contain extracellular bacteria released by dying cells (29). We observed that around 3% and 9% of Mm clusters are targeted by GFP-Lc3 at 1 and 2 dpi, respectively, which increases to uncountable levels at 3 dpi because of the increasing numbers and size of granulomas (Fig1 B and Fig1 C). These results were confirmed by Western blot, showing that LC3-II protein levels – indicative of autophagosome formation – gradually increased during Mm infection compared to uninfected controls (Fig1 D). Using a FK2 ubiquitin antibody, which can recognize monoubiquitinated cell surface molecules as well as polyubiquitin chains, we observed that ubiquitin co-colocalized with approximately 4% and 10% of the Mm clusters at 1 and 2 dpi, respectively (Fig1 E and Fig1 F). Furthermore, we observed by Western blot detection that Mm infection increased general levels of protein ubiquitination (Fig1 G). In addition, we found that ubiquitin and GFP-Lc3 co-localized at Mm clusters (Fig1 H). Collectively, these data demonstrate that Mm is marked by ubiquitin and that overall ubiquitination levels are induced during infection in the zebrafish model, which coincides with autophagic targeting of bacteria.

**Fig 1.**
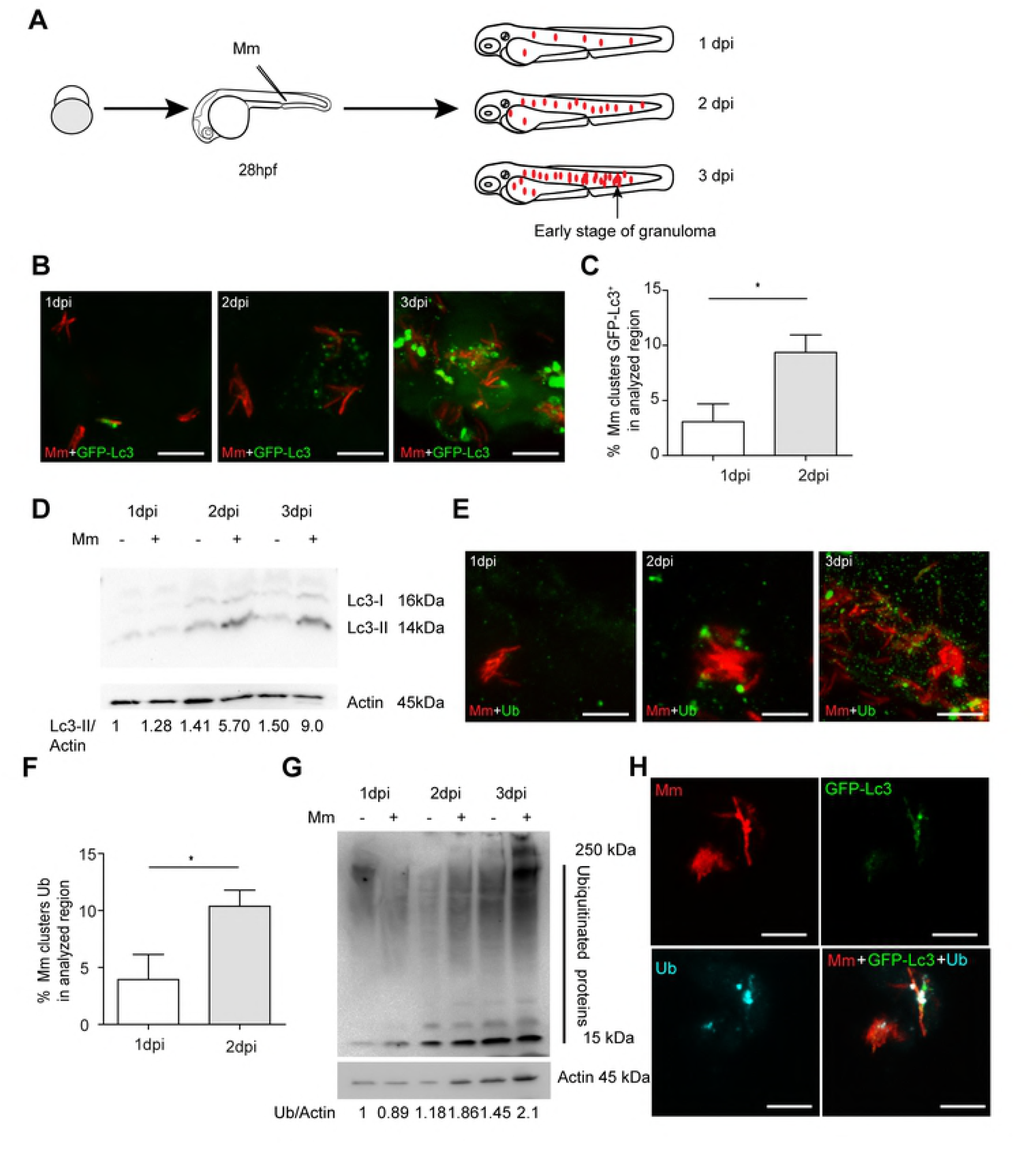
Ubiquitination and autophagy activity can be induced by Mm infection. (A) Schematic diagram of the zebrafish Mm infection model for TB study. *Mycobacterium marinum* (Mm) strain 20 fluorescently labelled with mCherry was microinjected into the blood island of embryos at 28 hpf. Red dots represent small clusters of Mm-infected cells visible from 1 dpi. At 3 dpi these Mm clusters have grown into early stage granulomas. (B) Representative confocal micrographs of GFP-Lc3 co-localization with Mm clusters in infected embryos/larvae at 1, 2 and 3 days post infection (dpi). Scale bars, 10 μm. (C) Quantification of the percentage of Mm clusters positive for GFP-Lc3 at 1 and 2 dpi. The results are representative for two individual repeats (≥ 20 embryo/group). ns, non-significant,*p<0.05,**p<0.01,***p<0.001. (D) Western blot determination of Lc3 protein levels in infected and uninfected embryos/larvae at 1, 2 and 3 dpi. Protein samples were extracted from 1, 2 and 3 dpi infected and uninfected larvae (>10 larvae/sample). The blots were probed with antibodies against Lc3 and Actin as a loading control. Western blot was representative for three independent experimental repeats. (E) Representative confocal micrographs of Ubiquitin co-localization with Mm clusters in infected embryos/larvae at 1, 2 and 3 days post infection (dpi). Scale bars, 10 μm. (F) Quantification of the percentage of Mm clusters positive for ubiquitin staining at 1 and 2 dpi (≥ 10 embryo/group). The results are representative for two individual repeats. ns, non-significant, *p<0.05, **p<0.01, ***p<0.001. (G) Western blot analysis of ubiquitination levels in infected and uninfected embryos/larvae at 1, 2 and 3 dpi. Protein samples were extracted from 1, 2 and 3 dpi infected and uninfected larvae (>10 larvae/sample). The blots were probed with an antibody detecting both poly and mono ubiquitin and with anti-Actin antibody as a loading control. Western blot representative for three independent experimental repeats. (H) Representative confocal micrographs of GFP-Lc3 and Ubiquitin co-localization with Mm clusters in infected larvae at 3 dpi. Scale bars, 10 μm.

### Deficiency in the ubiquitin receptors Optineurin or p62 does not impair zebrafish development

Since ubiquitinated bacteria are targets for members of the sequestosome-like receptor family, we compared the protein sequences of its members p62, Optineurin, Calcoco2 (Ndp52), Nbrc1, and Tax1bp1 between human, zebrafish and other vertebrates, showing a high overall degree of conservation (S1B Fig and S1C Fig). We focused this study on two members of the family, *p62* and *optn*, which are transcriptionally induced during Mm infection of zebrafish based on published RNA sequencing data (27) and show strong similarity with their human orthologues in the ubiquitin-binding domains (UBA in p62 and UBAN in Optineurin) and Lc3 interaction regions (LIR) (S1D Fig). With the aim to investigate the functions of Optineurin and p62 in anti-mycobacterial autophagy, we utilized CRISPR/Cas9 genome editing technology to generate mutant zebrafish lines. We designed short guide RNAs for target sites at the beginning of coding exons 2 of the *optn* and *p62* genes, upstream of the exons encoding the ubiquitin and Lc3 binding regions, such that the predicted effect of CRISPR mutation is a complete loss of protein function (Fig2 A). A mixture of sgRNA and Cas9 mRNA was injected into zebrafish embryos at the one cell stage and founders carrying the desired mutations were outcrossed to the *Tg*(*CMV:EGFP-map1lc3b*) autophagy reporter line (hereafter referred to as GFP-Lc3) (Fig2 B)(30). The established *optn* mutant allele carried a 5 nucleotides deletion at the target site, which we named *optn*^Δ5n/Δ5n^ (Fig2 C). The *p62* mutant allele carried an indel resulting in the net loss of 37 nucleotides, which we named *p62*^Δ37n/Δ37n^ (Fig2 C). The homozygous mutants were fertile and produced embryos that did not exhibit detectable morphological differences compared with embryos produced by their wild-type (*optn*^+/+^ or *p62*^+/+^) siblings (S1A Fig). Furthermore, no significant deviation from the Mendelian 1:2:1 ratio for +/+, +/- and -/- genotypes was observed when the offspring of heterozygous incrosses were sequenced at 3 months of age (Fig2 E). Western blot analysis using anti-Optineurin and anti-p62 C-terminal antibodies confirmed the absence of the proteins in the respective mutant lines (Fig2 D). In addition, quantitative PCR (Q-PCR) analysis revealed approximately 4.5-fold reduction of *optn* mRNA in the *optn*^Δ5n/Δ5n^ larvae and 10-fold reduction of *p62* mRNA in the *p62*^Δ37n/Δ37n^ larvae, indicative of nonsense-mediated mRNA decay (Fig2 F). Collectively, the *optn*^Δ5n/Δ5n^ and *p62*^Δ37n/Δ37n^ mutant zebrafish produce no functional Optineurin or p62, respectively, and the loss of these ubiquitin receptors does not induce detectable developmental defects that could interfere with the use of the mutant lines in infection models.

**Fig 2.**
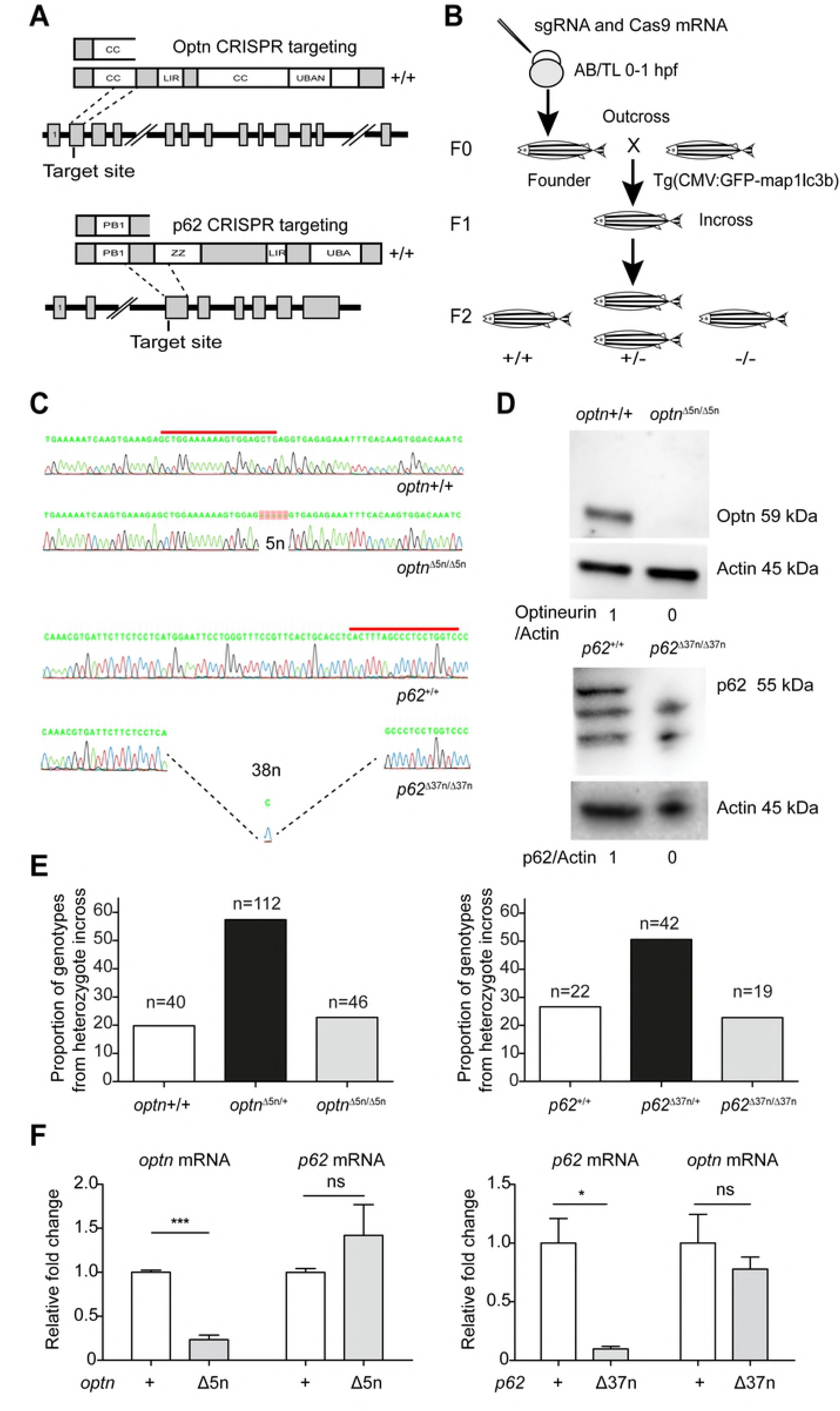
Generation of Optineurin and p62 mutant lines. (A) Schematic representation of the Optn and p62 genetic and protein domain architecture and CRISPR target site. Optn (517 aa) and p62 (452 aa) both contain a Lc3 interaction region domain (LIR) and ubiquitin binding domains (UBAN in Optn and UBA in p62). Additionally, two coiled-coil motifs (CC) in Optineurin and the PHOX/Bem1p (PB) and Zinc Finger (ZZ) domains of p62 are indicated. The gene loci are shown with coding exons as grey boxes (14 in Optn and 8 in p62) and introns as solid black lines (large introns not drawn to scale). The position of the CRISPR target site sequences at the beginning of exon 2 in Optineurin and exon 3 in p62 are indicated and the predicted truncated proteins in the mutant lines are drawn above. (B) Schematic diagram of the generation of Optn and P62 mutant lines. Target-specific sgRNA and Cas9 mRNAs were co-injected into one cell stage embryos (AB/TL WT line). Founders were outcrossed to *Tg*(*CMV*:*EGFP-Lc3*) fish and the F1 was incrossed to obtain homozygous mutant and wild type F2 siblings. (C) Sanger sequencing of WT and mutant F2 fish. Red lines indicate CRISPR target sites. The Optn and p62 mutant sequences contain deletions (indels) of 5 and 37 nucleotides, respectively. (D) Confirmation of CRISPR mutation effect by Western blot analysis. Protein samples were extracted from 4 dpf *optn* or 3dpf *p62* mutant and WT larvae (>10 embryos/sample) and Western blots were repeated at least three times with independent extracts. The blots were probed with antibodies against Optn or P62 and Actin as a loading control. Optn/Actin and P62/Actin ratios) are indicated below. kDa, kilodalton. (E) Segregation from F1 heterozygous incross. Genotypes of adult fish (>3 months) combined from 4 (for *optn*) or 3 (*p62*) independent breedings were confirmed by PCR and sequencing. (F) *optn* and *p62* mRNA was detected by quantitative PCR. Total RNA was isolated from 4dpf of *optn*^+/+^, *optn*^Δ5n/Δ5n^, *p62*^+/+^ and *p62*^Δ37n/Δ37n^ embryos (>10 embryos/sample) from three biological replicates.

### Optineurin or p62 deficiencies affect autophagy

To analyze the effects of Optineurin or p62 deficiency on autophagy, we performed Lc3 Western blot detection on whole embryo extracts and imaged GFP-Lc3 signal *in vivo* (Fig3 A). Differences in the levels of the cytosolic (Lc3-I) and membrane-bound (Lc3-II) forms of Lc3 or effects on GFP-Lc3 puncta accumulation can be due to altered basal autophagy levels, but can also be caused by differences in autophagosome degradation. Therefore, we also examined Lc3-I/Lc3-II levels and GFP-Lc3 accumulation in larvae following treatment with Bafilomycin A1 (Baf A1), which is an inhibitor of vacuolar H+ ATPase (V-ATPase) that prevents maturation of autophagic vacuoles by inhibiting fusion between autophagosomes and lysosomes (31, 32). First, we performed a dose range assay to determine the effect of Baf A1 on Lc3-II accumulation in zebrafish embryos. Results showed that after 12 h of incubation, a dosage of 100nM resulted in Lc3-II accumulation without affecting the Lc3-I level, whereas higher dosage additionally increased the Lc3-I level (S2A Fig). Thus, we utilized a dosage of 100nM to test Lc3-II accumulation in wildtype and mutant embryos not carrying the GFP-Lc3 reporter (Fig3 B). No differences in Lc3-II accumulation were observed between *optn*^+/+^ and *optn*^Δ5n/Δ5n^ embryos or between *p62*^+/+^ and *p62*^Δ37n/Δ37n^ embryos (Fig3 C). However, accumulation of Lc3-II in *optn* or *p62* mutant embryos was significantly reduced in presence of Baf A1 (52% and 66%, respectively) compared to the wildtype controls (Fig3 C). In agreement, the number of GFP-Lc3 puncta in *optn* or *p62* mutants were significantly lower than in the corresponding WT controls, showing 59% and 47% reductions, respectively (Fig3 D and Fig3 E).

**Fig 3.**
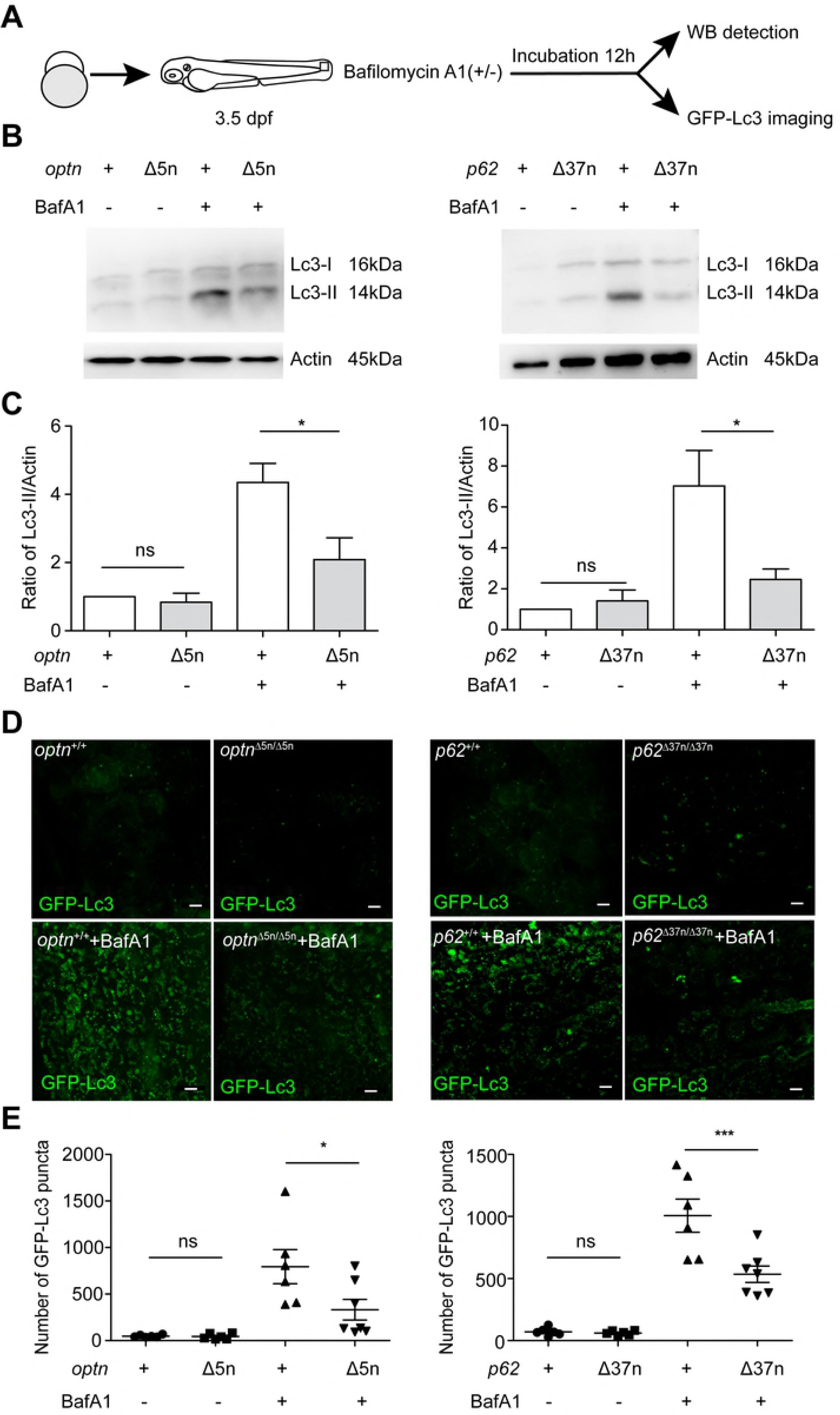
Optineurin or p62 deficiency affects autophagosome formation. (A) Workflow of the experiments shown in (B-G). 3.5 dpf larvae were treated with Bafilomycin A1 (Baf A1) (100 nM) for 12h. The GPF-Lc3 negative larvae were selected to assay autophagy activity by Western blot, the GFP-Lc3 positive larvae were collected to monitor autophagic activity using confocal imaging. The red square indicates the region for confocal imaging. (B) The level of basal autophagy in WT and mutant embryos in absence or presence of Baf A1. Protein samples were extracted from 4 dpf WT and mutant larvae (>10 embryos/sample). The blots were probed with antibodies against Lc3 and Actin as a loading control. Western blots were repeated at least three times with independent extracts. (C) Quantification of Lc3-II fold changes in WT and mutant embryos in absence or presence of Baf A1. Western blot band intensities were quantified by Lab Image. Data is combined from three independent experiments. (D) Representative confocal micrographs of GFP-Lc3 puncta present in the tail fin of *optn*^+/+^, *optn*^Δ5n/Δ5n^, *p62*^+/+^ and *p62*^Δ37n/Δ37n^ at 4 dpf . Scale bars, 10 μm. E. Quantification of the number of GFP-Lc3 puncta in *optn*^+/+^, *optn*^Δ5n/Δ5n^, *p62*^+/+^ and *p62*^Δ37n/Δ37n^ larvae with and without Baf A1 treatment. Each larva was imaged at a pre-defined region of the tail fin (as indicated by the red boxed area in Fig3 A) (≥6 larvae/group). Results are representative of two independent experiments.

The function of Optineurin and p62 as ubiquitin receptors implies that these proteins are degraded themselves during the process of autophagy. Therefore, we asked if p62 protein levels are affected in *optn* mutants or, vice versa, if *p62* mutation impacts Optineurin protein levels. Western blot analysis showed accumulation of p62 and Optineurin protein in wild type embryos in response to Baf A1 treatment, confirming that these ubiquitin receptors are substrates for autophagy under basal conditions (S2B Fig). Levels of p62 protein were reduced in *optn*^Δ5n/Δ5n^ embryos compared with *optn*^+/+^, both in absence or presence of Baf A1 (Fig3 F). This difference was not due to a transcriptional effect, since p62 mRNA levels were not significantly different between *optn*^+/+^ and *optn*^Δ5n/Δ5n^ embryos (Fig2 F). Similarly, levels of Optineurin protein were reduced in *p62*^Δ37n/Δ37n^ embryos compared with *p62*^+/+^ in absence or presence of Baf A1 (Fig3 F), and again this was not associated with a difference in mRNA expression (Fig2 F). In conclusion, the absence of either of the ubiquitin receptors, Optineurin or p62, leads to increased use of the other ubiquitin receptor as a substrate for autophagic degradation. Furthermore, loss of either of the receptors leads to lower levels of Lc3-II and GFP-Lc3 accumulation when lysosomal degradation is blocked, suggesting reduced activity of the autophagy pathway in the *optn* and *p62* mutants.

### Optineurin or p62 deficiencies increase the susceptibility of zebrafish embryos to Mm infection

Next, we asked if *optn* or *p62* mutations would affect the resistance of zebrafish embryos to mycobacterial infection. We injected Mm into embryos via the caudal vein at 28 hpf to measure infection burden at 3 dpi (Fig4 A). The infection data showed that *optn* or *p62* mutant embryos were hypersusceptible to Mm infection compared with their WT controls, culminating in an increase of the Mm fluorescent signal of 2.8 and 2.9 times, respectively (Fig4 B). In addition, we examined whether transient knockdown of *optn* or *p62* would phenocopy the infection phenotype of the mutant lines. We injected *optn* or *p62* antisense morpholino oligonucleotides into the one cell stage of embryos and collected injected individuals at 28h for confirmation of the knockdown effect by reverse transcription polymerase chain reaction (RT-PCR) and Western blot (S3A Fig, S3B Fig and S3C Fig). Subsequently, analysis of the Mm infection burden at 3 dpi showed that transient knockdown of *optn* or *p62* led to similar increases of the Mm infection burden as had been observed in the mutant lines (Fig4 C). Since Optineurin and p62 are known to function cooperatively in xenophagy of *Salmonella enterica* (22-24), we asked if double deficiency of Optineurin and p62 resulted in an increased infection burden compared to single mutation of either *optn* or *p62*. No additive effect on the infection burden was observed when *p62* morpholino was injected into *optn* mutant embryos or *optn* morpholino into *p62* mutant embryos (Fig4 D). Taken together, our data demonstrate that both Optineurin and p62 are required for controlling Mm infection and that loss of either of these ubiquitin receptors cannot be compensated for by the other receptor in this context.

**Fig 4.**
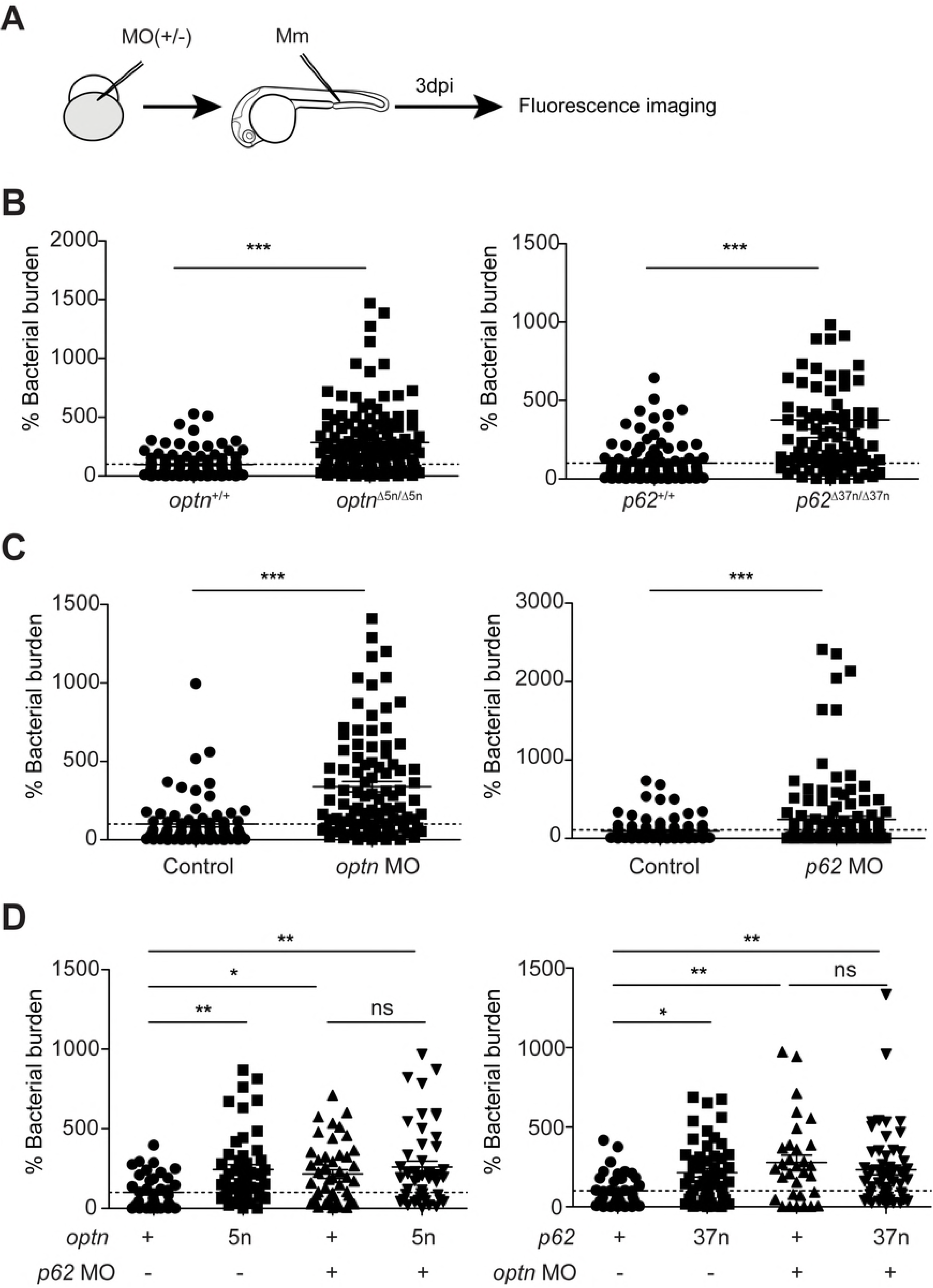
Optineurin or p62 deficiency leads to increased susceptibility to Mm infection. (A) Workflow of the experiments shown in (B-D). *optn* or *p62* MO were injected into the one cell stage of embryos and infection was performed at 28 hpf with 200 CFU of Mm via blood island microinjection. Bacterial quantification was done at 3dpi. (B-D) Mm infection burden in *optn* and *p62* mutant larvae (B), under *optn* and *p62* MO knockdown conditions (C), and following injection of *p62* MO or *optn* MO in *optn* and *p62* mutants, respectively (D). The data are accumulated from three independent infection experiments. Each dot represents an individual larva. ns, non-significant,*p<0.05,**p<0.01,***p<0.001.

### Optineurin or p62 deficiency reduces the autophagy response to Mm infection

Having established that mutation of either *optn* or *p62* results in increased Mm infection burden, we investigated if the inability of mutant embryos to control infection is due to a reduction in the targeting of mycobacteria to autophagy (Fig5 A). To this end, we first examined the association of GFP-Lc3 with Mm at 1 dpi. Mm has formed small infection foci at this time point, which could be manually scored as positive or negative for GFP-Lc3 association. In wild type embryos 5-6% of these infection foci were positive for GFP-Lc3 (S4A Fig and S4B Fig). The percentage of GFP-Lc3 positive Mm clusters was approximately 50% lower in the *optn* or *p62* mutant embryos compared with their wild type controls, but differences were not statistically significant due to the relatively low number of these GFP-Lc3 association events (S4A Fig and S4B Fig). We continued to examine GFP-Lc3 targeting to Mm at 2 dpi and found that mutation of *optn* or *p62* resulted in significantly decreased GFP-Lc3 co-localization with Mm clusters (Fig5 A, B and C). In addition, we used GFP-Lc3-negative mutant and wild type larvae for Western blot analysis of Lc3-II protein levels in response to infection. We found that Mm infection increased Lc3-II protein levels approximately 3- to 5-fold in wild type (*optn*^+/+^ amd *p62*^+/+^) larvae at 3 dpi, whereas this induction level was approximately 50% lower in the *optn* and *p62* mutant larvae (Fig5 D). Mm-infected mutant embryos also showed reduced Lc3-II accumulation in the presence of Baf A1 (S4B Fig). Taken together, these data support the hypothesis that Optineurin and p62 are required for autophagic defense against mycobacterial infection.

**Fig 5.**
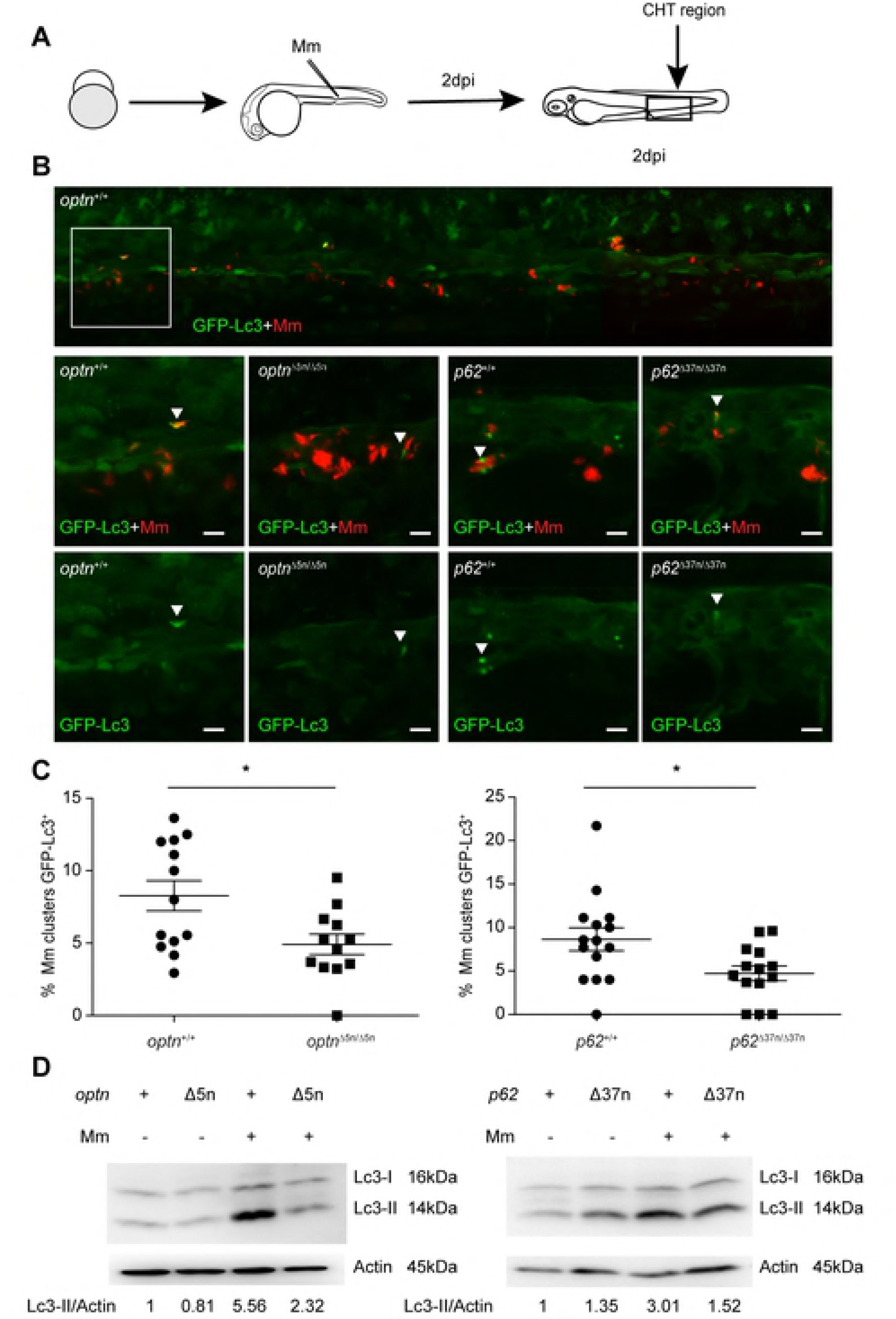
Optineurin or p62 deficiency inhibits targeting of Mm by GFP-Lc3. (A) Workflow of the experiment shown in B. 2 dpi fixed larvae were used for confocal imaging. The entire caudal hematopoietic tissue (CHT) was imaged, as indicated by the black box. (B) Representative confocal micrographs of GFP-Lc3 co-localization with Mm clusters in infected larvae. The top image shows the entire CHT region in *optn*^+/+^ infected larvae. The area indicated by the white box is detailed below. The bottom images show GFP-Lc3 co-localization of Mm clusters in *optn*^+/+^, *optn*^Δ5n/Δ5n^, *p62*^+/+^ and *p62*^Δ37n/Δ37n^ infected larvae. The arrowheads indicate the overlap between GFP-Lc3 and Mm clusters. Scale bars, 10 μm. (C) Quantification of the percentage of Mm clusters positive for GFP-Lc3 vesicles. The data is accumulated from two independent experiments; each dot represents an individual larva (≥12 larvae/group). ns, non-significant, *p<0.05,**p<0.01,***p<0.001. (D) Lc3 protein levels were determined by Western blot in infected and uninfected larvae. Protein samples were extracted from 4 dpf larvae (>10 larvae/sample). The blots were probed with antibodies against Lc3 and Actin as a loading control. Western blots were repeated two times with independent extracts.

### Overexpression of *optn* or *p62* increases resistance of zebrafish embryos to Mm infection

To further test the hypothesis that Optineurin and p62 mediate autophagic defense against Mm, we generated full-length *optn* and *p62* mRNAs *in vitro* and injected these into embryos at the one cell stage, resulting in ubiquitous overexpression (Fig6 A). The increase in Optineurin or p62 protein levels following mRNA injection was verified by Western blot analysis (Fig6 B) and no effects of overexpression on embryo survival or development were observed (data not shown). Overexpression of *optn* or *p62* mRNAs significantly reduced Mm infection burden at 2 or 3 dpi compared to the control groups (Fig6 C and S5A Fig). Furthermore, injection of *optn* or *p62* mRNAs carrying deletions in the sequences encoding the ubiquitin binding domains or Lc3 interaction regions did not lead to a reduction of the Mm infection burden compared with the control groups (Fig6 C). Thus, we conclude that *optn* or *p62* overexpression protects against Mm infection in a manner dependent on the interaction of the Optn and p62 proteins with both ubiquitin and Lc3.

**Fig 6.**
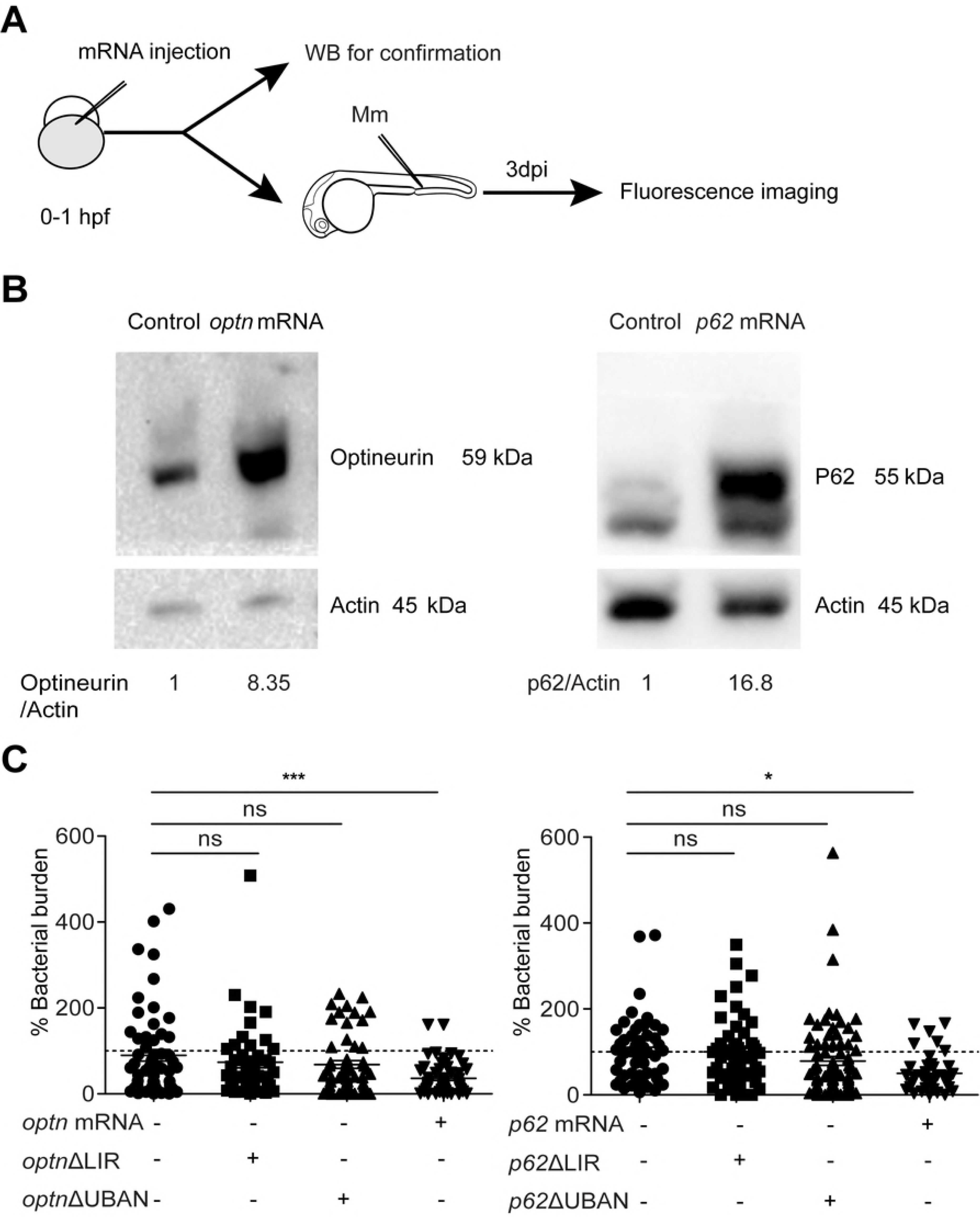
Transient overexpression *optn* or *p62* mRNA protects against Mm infection. (A) Workflow representing the experimental design in (B-C). *optn* or *p62* mRNA was injected into the one cell stage of embryos (*AB/TL*) at a dosage of 100 pg/embryo. Injected embryos were collected at 28 hpf for confirmation of the overexpression by Western blot analysis. Embryos were infected at 28hpf with 200 CFU Mm via the blood island by microinjection and bacterial burden was determined at 3 dpi. (B) Western blot analysis to test the effect of transient overexpression of *optn* or *p62* mRNA. Protein extracts were made from >20 mRNA-injected or control embryos per group. The blots were probed with antibodies against Optineurin or p62 and Actin as a loading control. Similar results were observed in two independent experiments. (C) Quantification of Mm infection burden in embryos injected with full length or ΔLIR/ΔUBAN deletion mRNAs of *optn* and *p62*. Accumulated data from two independent infection experiments is shown. ns, non-significant,*p<0.05,**P<0.01,***p<0.001.

### Overexpression of *optn* or *p62* promotes GFP-Lc3 association with Mm

Since overexpression of *optn* or *p62* mRNAs resulted in decreased Mm infection burden, we postulated that elevation of the Optn or p62 protein levels would result in increased targeting of Mm to autophagy by these ubiquitin receptors, in a manner dependent on the functions of the Lc3 interaction (LIR) and ubiquitin binding domains (UBAN/UBA). To test this hypothesis, we injected the full-length mRNAs, or mRNAs generated from deletion constructs lacking these domains, and quantified GFP-Lc3-positive and GFP-negative Mm infection foci at 1 dpi and 2 dpi (S6A Fig and Fig7 A). The results showed that overexpression of full-length *optn* or *p62* mRNAs significantly increased the percentage of GFP-Lc3-positive Mm clusters at 2 dpi, compared with the control groups (Fig7 B and Fig7 C). Conversely, injection of *optn* ΔUBAN, *optn* ΔLIR, *p62* ΔUBA and *p62* ΔLIR mRNAs did not increase the association of GFP-Lc3 with Mm clusters (Fig7 B and Fig7 C). Similar results could be observed as early as 1 day post infection (S6B Fig). In conclusion, our combined results demonstrate that Optineurin and p62 can target Lc3 to Mm and that increasing the level of either of these receptors promotes host defense against this mycobacterial pathogen.

**Fig 7.**
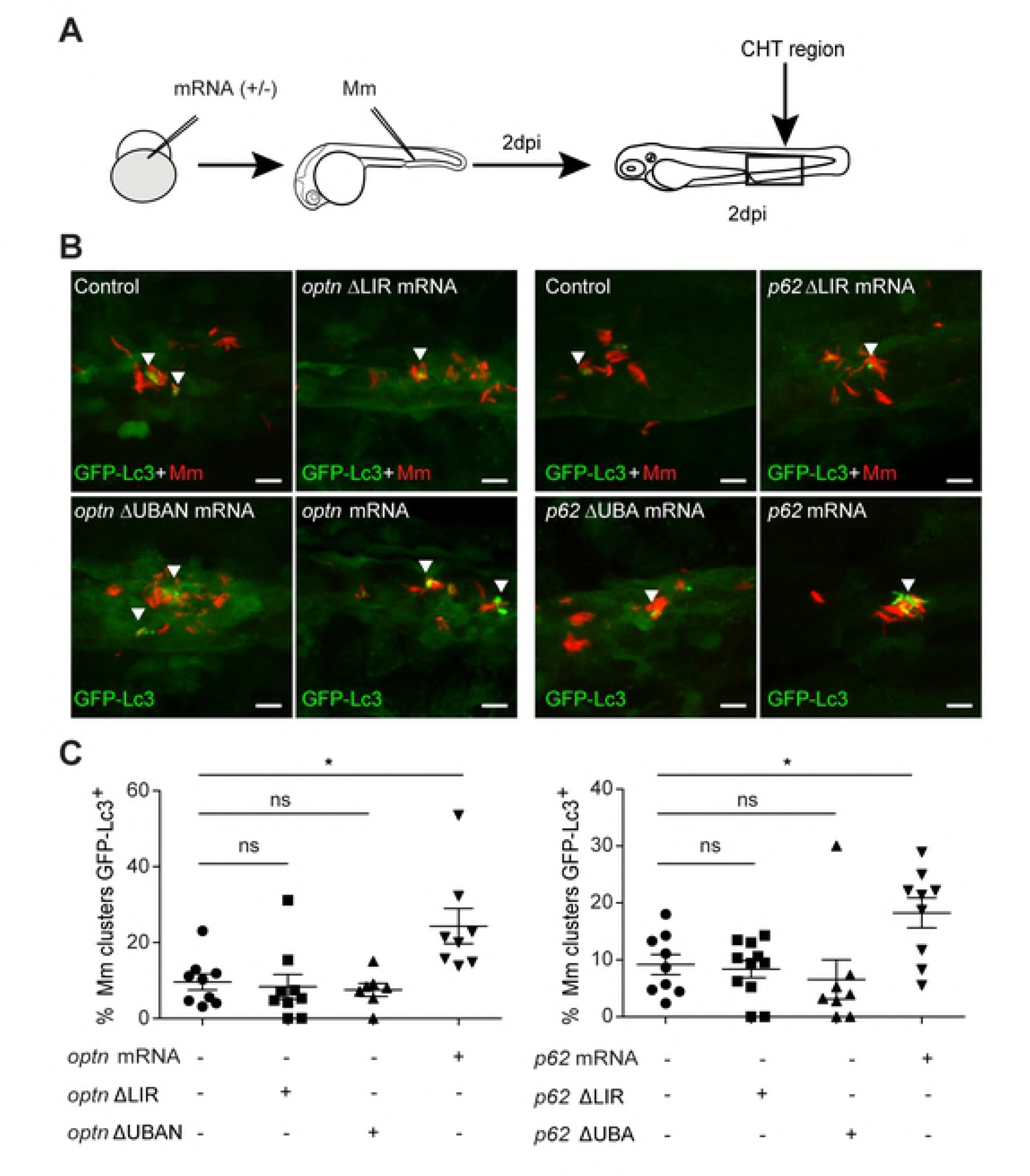
Transient overexpression of *optn* or *p62* mRNA promotes GFP-Lc3 recruitment to Mm clusters. (A) Workflow of the experiments in (B-C). *optn* or *p62* mRNA was injected into the one cell stage of embryos at a dosage of 100 pg/embryo. 2 dpi fixed larvae were used for confocal imaging. The entire caudal hematopoietic tissue (CHT) was imaged, as indicated by the black box. (B) Representative confocal micrographs of GFP-Lc3 co-localization with Mm clusters in larvae injected with full length or ΔLIR/ΔUBAN deletion mRNAs of *optn* and *p62*. The arrowheads indicate the overlap between GFP-Lc3 and Mm clusters. Scale bars, 10 μm. (C) Quantification of the percentage of Mm clusters positive for GFP-Lc3 vesicles. Each dot represents an individual larva (≥7 larvae/group). Results are representative of two independent experiments. ns, non-significant,*p<0.05,**P<0.01, *** p<0.001.

## Discussion

Members of the family of sequestosome (p62/SQSTM1)-like receptors (SLRs) function in autophagic host defense mechanisms targeting a range of intracellular pathogens, including *Salmonella, Shigella*, *Streptococci*, *Listeria*, *Mycobacteria*, and Sindbis virus (5, 13, 14, 33). These discoveries inspired investigations into autophagy modulators as host-directed therapeutics for treatment of infectious diseases, including Tb (9, 34, 35). However, the relevance of autophagic defense mechanisms for host resistance against Mtb infection has recently been questioned (15, 36). This indicates that there are significant gaps in our understanding of the interaction between components of the autophagy pathway and mycobacterial pathogens, emphasizing the need for more research in animal models of Tb (12). Here, we have studied the function of two SLR family members in the zebrafish Tb model. We show that selective autophagy mediated by p62 and Optineurin provides resistance against mycobacterial infection in the context of our *in vivo* infection model that is representative of the early stages of Tb granuloma formation (17, 19). Our findings support the host-protective role of p62 in Tb by autophagic targeting of *Mycobacteria*, in line with previous *in vitro* studies (13, 14). Importantly, we also present the first evidence linking Optineurin to resistance against *Mycobacteria*, expanding our understanding of the function of SLRs in host defense against intracellular pathogens.

The zebrafish embryo and larval Tb model provides the opportunity to image critical stages of the mycobacterial infection process, from the initial phagocytosis of Mm by macrophages up to the early stages of Tb granuloma formation (37). The model is representative of miliary Tb, where the infection is disseminated to multiple organs of the host. The embryonic and larval stages of the zebrafish allow us to study the contribution of innate immunity to host defense, since they lack a matured adaptive immune response at this time point of development (17). We therefore used this model to study the importance of autophagic defense mechanisms during innate host defense against mycobacterial infections. In this study, we successfully generated *p62* and *optn* loss-of-function zebrafish mutant lines using CRISPR/Cas9 technology. Besides its role in host defense, p62 is a stress-inducible protein that functions as a signalling hub in diverse processes like amino acid sensing and the oxidative stress response (38). Defects in autophagy pathways caused by mutations in *OPTN* have been associated with human disorders like glaucoma, Paget disease of bone, and amyotrophic lateral sclerosis (24, 39). Despite the important functions reported for p62 and Optineurin in cellular homeostasis, the mutant fish lines we generated are viable and fertile. The absence of either p62 or Optineurin resulted in increased use of the other ubiquitin receptor to sequester autophagic cargo in zebrafish larvae. Nonetheless, loss of either of the receptors leads to lower levels of Lc3-II and GFP-Lc3 accumulation when lysosomal degradation is blocked, which indicates reduced activity of the autophagy pathway in these mutants. Therefore, we could use these mutant lines to gain a better understanding of the role of p62, Optineurin, and selective autophagy in host defense against mycobacterial infection.

Genetic links between autophagy pathway genes and susceptibility to Tb in human populations support the function of autophagy in innate host defense against Mtb (40). However, the contribution of autophagy as a direct anti-mycobacterial mechanism has recently been challenged, since macrophage-specific depletion of a number of autophagy genes, including *p62*, did not affect the outcome of disease in a mouse model of Tb (15, 36). A possible explanation for these findings, as suggested by the authors of this study, is that Mtb, like other successful intracellular pathogens, could have evolved virulence mechanisms that subvert or exploit autophagic defense mechanisms employed by the host (41). In case of one of the autophagy genes, *ATG5*, macrophage-specific depletion increased Mtb infection in mice by over-activating inflammation rather than by impairing autophagic processes (15). It is therefore conceivable that modulating the activity of SLRs could also affect inflammation. Indeed, Optineurin has been implicated in inflammatory bowel disease and both p62 and Optineurin are involved in regulation of inflammatory signaling downstream of NF-κB (42-46). Through a process that involves polyubiquitination of regulatory proteins, both p62 and Optineurin can modulate the activity of the IKK kinase complex that activates NFκB (42, 43). It is therefore possible that altered inflammatory responses in *p62* and *optn* mutants could explain (part of) the increase in mycobacterial burden observed in zebrafish hosts, while the beneficial role for autophagic defense mechanisms targeting the bacteria might be limited.

To investigate the possible role of Optineurin and p62 in anti-mycobacterial autophagy, we quantified the association between GFP-Lc3 and Mm under loss-of-function and gain-of-function conditions of both receptors. In wild type zebrafish embryos, only 3-5% of the bacteria co-localized with autophagic vesicles one day after a systemic infection with mycobacteria. Although the number of GFP-Lc3 positive bacterial clusters rises over the next two days, the percentage of bacteria targeted by autophagy at any distinct time point remains relatively low (e.g. ∼10% at 2 days post infection). According to these results, the host only employs autophagic defense mechanisms against a small proportion of the invading mycobacteria during early stages of the infection, either because there is no greater need, or because the pathogens are indeed effectively suppressing this response. It is important to note though that GFP-Lc3 association with Mm is a transient process (20), which means that the percentage of bacteria that encounter autophagic defenses throughout the early infection process might be much higher. Strikingly, the percentage of bacteria labeled by ubiquitin closely resembled the percentage of bacteria targeted by autophagy, and we were able to detect clear colocalization between ubiquitin and GFP-Lc3 at bacterial clusters. Upon loss-of-function of either p62 or Optineurin, the co-localization between bacteria and autophagic vesicles decreased and the bacterial burden increased. Conversely, overexpression of either ubiquitin binding receptor increased autophagic targeting of bacteria and resulted in lower bacterial burdens, both of which required the presence of functional Lc3 and ubiquitin binding domains. Taken together, we conclude that autophagic targeting of mycobacteria by p62 and Optineurin indeed provides protection against infection in our *in vivo* Tb model.

In summary, our findings confirm that p62 mediates ubiquitin-dependent autophagic targeting of mycobacteria in an *in vivo* model for Tb. We also provide the first evidence that the SLR family member Optineurin is involved in autophagic targeting of ubiquitinated mycobacteria. While we cannot exclude a role for p62 and Optineurin in regulating inflammatory processes during Tb disease progression, we have shown that the autophagic targeting of mycobacteria by these ubiquitin-binding receptors forms an important aspect of innate host defense against Tb. Our results are therefore especially important for the development of new treatment strategies for Tb patients with a compromised adaptive immune system – such as in HIV-coinfection. Based on these results, selective autophagy stimulation remains a promising strategy for development of novel anti-Tb therapeutics.

## Materials and methods

### Ethics statement

Zebrafish lines in this study (S1 Table) were handled in compliance with local animal welfare regulations as overseen by the Animal Welfare Body of Leiden University (License number: 10612) and maintained according to standard protocols (zfin.org). All protocols adhered to the international guidelines specified by the EU Animal Protection Directive 2010/63/EU. The generation of zebrafish *optn* and *p62* mutant lines was approved by the Animal Experimention Committee of Leiden University (UDEC) under protocol 14198. All experiments with these zebrafish lines were done on embryos or larvae up to 5 days post fertilization, which have not yet reached the free-feeding stage. Embryos were grown at 28.5°C and kept under anesthesia with egg water containing 0.02% buffered 3-aminobenzoic acid ethyl ester (Tricaine, Sigma) during bacterial injections, imaging and fixation.

### CRISPR/Cas9 mediated mutagenesis of zebrafish *optn* and *p62*

Single guide RNAs (sgRNAs) targeting the second coding exon of zebrafish *optn* (ENSDART00000014036.10) and the third coding exon of *p62 (*ENSDART00000140061.2) were designed using the chop-chop website (47). To make sgRNAs, the template single strand DNA (ssDNA) (122 bases) was obtained by PCR complementation and amplification of full length ssDNA oligonucleotides. Oligonucleotides up to 81 nucleotides were purchased from Sigma-Aldrich using standard synthesis procedures (25 nmol concentration, purification with desalting method) (S2 Table and S3 Table). The pairs of semi-complimentary oligos were annealed together by a short PCR program (50 µL reaction, 200uM dTNPs, 1 unit of Dream Taq polymerase (EP0703, ThermoFisher); PCR program: initial denaturation 95°C/3 minute (min), 5 amplification cycles 95°C/30 Second (s), 55°C/60 s, 72°C/30 s, final extension step 72°C/15 min) and subsequently the products were amplified using the primers in S2 Table with a standard PCR program (initial denaturation 95°C/3 min, 35 amplification cycles 95°C/30 s,55°C/60 s, 72°C/30 s, final extension step 72°C/15 min). The final PCR products were purified with Quick gel extraction and PCR purification combo kit (00505495, ThermoFisher). The purified PCR products were confirmed by gel electrophoresis and Sanger sequencing (Base Clear, Netherlands). For *in vitro* transcription of sgRNAs, 0.2 µg template DNA was used to generate sgRNAs using the MEGA short script ®T7 kit (AM1354, ThermoFisher) and purified by RNeasy Mini Elute Clean up kit (74204, QIAGEN Benelux B.V., Venlo, Netherlands). The Cas9 mRNA was transcribed using mMACHINE® SP6 Transcription Kit (AM1340, Thermo Fisher) from a Cas9 plasmid (39312, Addgene) (Hrucha et al 2013) and purified with RNeasy Mini Elute Clean up kit (74204,QIAGEN Benelux B.V., Venlo, Netherlands). A mixture of sgRNA and Cas9 mRNA was injected into one cell stage AB/TL embryos (sgRNA 150 pg/embryo and Cas9 mRNA 300 pg/embryo). The effect of CRISPR injection was confirmed by PCR and Sanger sequencing.

### Genomic DNA isolation and genotyping

Genomic DNA was isolated from an individual embryo (2 dpf) or small pieces of the tail fin tissue of adults (>3 months) by fin clipping. Embryos or tissue samples were incubated in 200 µL 100% Methanol at −20°C overnight (O/N), then methanol was removed, and remaining methanol was evaporated at 70°C for 20 min. Next, samples were incubated in 25 µL of TE buffer containing 1.7 µg/µL proteinase K at 55°C for more than 5 h. Proteinase K was heat inactivated at 80°C for 30 min, after which samples were diluted with 100 µL of Milli-Q water. Genotyping was performed by PCR-amplification of the region of interest using the primers in S5 Table followed by Sanger sequencing to identify mutations (Base Clear, Netherlands).

### Western blot analysis

Embryos (28hpf/2dpf/4dpf/3dpi) were anaesthetised with Tricaine (Lot#MKBG4400V, SIGMA-ALDRICH) and homogenised with a Bullet-blender (Next-Advance) in RIPA buffer (#9806, Cell Signalling) containing a protein inhibitor cocktail (000000011836153001, cOmplete, Roche). The extracts were then spun down at 4°C for 10 min at 12000 rpm/min and the supernatants were frozen for storage at −80°C. Western blot was performed using Mini-PROTEAN-TGX (456-9036, Bio-Rad) or 18% Tris—Hcl 18% polyacrylamide gels, and protein transfer to commercial PVDF membranes (Trans-Blot Turbo-Transfer pack, 1704156, Bio-Rad). Membranes were blocked with 5% dry milk (ELK, Campina) in Tris buffered saline (TBS) solution with Tween 20 (TBST, 1XTBS contains 0.1% Tween 20) buffer and incubated with primary and secondary antibodies. Digital images were acquired using Bio-Rad Universal Hood II imaging system (720BR/01565 UAS). Band intensities were quantified by densitometric analysis using Image Lab Software (Bio-Rad, USA) and values were normalised to actin as a loading control. Antibodies used were as follows: polyclonal rabbit anti-Optineurin (C-terminal) (1:200, lot#100000; Cayman Chemical), polyclonal rabbit anti-p62 (C-terminal) (PM045, lot#019, MBL), polyclonal rabbit anti Lc3 (1:1000, NB100-2331, lot#AB-3, Novus Biologicals), Anti mono-and polyubiquitinated conjugates mouse monoclonal antibody (1:200; BML-PW8810-0100, lot#01031445, Enzo life Sciences), Polyclonal actin antibody (1:1000, 4968S, lot#3, Cell Signaling), Anti-rabbit IgG, HRP-Linked Antibody (1:1000, 7074S, Lot#0026, Cell Signaling), Anti-mouse IgG, HRP-linked Antibody (1:3000, 7076S, Lot#029, Cell Signaling).

### Morpholino design and validation

*optn* and *p62* splice blocking morpholinos were purchased from Gene Tools. For morpholino sequences see S4 Table. Morpholinos were diluted in Milli Q water with 0.05% phenol red and 1 nL of 0.1 mM *optn* or 0.5 mM p62 Morpholino was injected into the one cell stage of embryos as previously described (21). The knockdown effect was validated by RT-PCR and Western blot.

### Infection conditions and bacterial burden quantification

*Mycobacterium marinum* strain 20 bacteria, fluorescently labelled with mCherry, were microinjected into the blood island of embryos at 28 hpf as previously described (48). The injection dose was 200 CFU for all experiments. Before the injection, embryos were manually dechorionated around 24hpf. Approximately 5 min before bacterial injections, zebrafish embryos were brought under anaesthesia with tricaine. Infected embryos were imaged using a Leica MZ16FA stereo fluorescence microscopy with DFC420C camera, total fluorescent bacterial pixels per infected fish were determined on whole-embryo stereo fluorescent micrographs using previously described software (49).

### Confocal laser scanning microscopy and image quantification

Fixed or live embryos were mounted with 1.5% low melting agarose (140727, SERVA) and imaged using a Leica TCS SPE confocal microscope. For quantification of basal autophagy, fixed uninfected 4dpf larvae were imaged by confocal microscopy with a 63x water immersion objective (NA 1.2) in a pre-defined region of the tail fin to detect GFP-LC3-positive vesicles (Fig3 D and Fig3 E). The number of GFP-Lc3 vesicles per condition was quantified using Fiji/ImageJ software (Fig3 D and Fig3 E). For quantification of the autophagic response targeted to Mm clusters (Fig1 B and C, S4A Fig and B, S6A Fig and B), live or fixed infected embryos were viewed by confocal microscopy with a 63x water immersion objective (NA 1.2) and the number of Mm clusters that were targeted by GFP-Lc3 puncta in the tail region were counted manually. The same approach was used to quantify Ubiquitin targeting to Mm clusters (Fig1 E and F). To quantify the percentage of GFP-Lc3^+^ Mm clusters, we imaged the entire caudal hematopoietic tissue (CHT) region of 2 dpi larvae (confocal microscopy; 40X water immersion objective with NA 1.0) and stitched multiple images together to manually count the number of Mm clusters positive for GFP-Lc3 out of the total number of clusters (Fig5 B and C, Fig7 B and C)

### Immunostaining

Embryos (1,2,3 dpi) were fixed with 4% PFA in PBS and incubated overnight with shaking at 4°C. After washing the embryos three times briefly in PBS with 0.8% Triton-x100) (PBSTx), the embryos/larvae were digested in 10 µg/ml proteinase K (000000003115879001, SIGMA-ALDRICH) for 10 minutes at 37°C. Subsequently, the embryos were quickly washed, blocked with PBSTx containing 1% Bovine serum albumins (BSA) (A4503-100g, SIGMA-ALDRICH) for 2h at room temperature and incubated overnight at 4°C in mono-and polyubiquitinated conjugates mouse monoclonal antibody (1:200; BML-PW8810-0100; Enzo lifes Siences), diluted in the blocking buffer. Next, embryos were washed three times in PBSTx, incubated for 1 h in blocking buffer at room temperature, incubated for 2 h at room temperature in 1:200 dilution of Alexa Fluor 488 or 633 goat anti-mouse (Invitrogen) in blocking buffer, followed with three times washes in PBSTx for imaging.

### mRNA preparation and injection

*optn* (ENSDART00000014036.10, Ensembl) and *p62* (ENSDART00000140061.2, Ensembl) cDNAs were amplified from 3dpf AB/TL embryos by PCR (primers in S5 Table) and ligated into a vector using the Zero-blunt cloning PCR kit (450245, Invitrogen). The sequence was confirmed by Sanger sequencing (BaseClear, Netherlands), after which *optn* and *p62* cDNAs were subcloned into a pCS2+ expression vector.

*optn* ΔUBAN cDNA was produced by in vitro transcription of *optn*–pCS2+ constructs digested by Sca1(R3122, NEB), which excludes the region encoding the UBAN protein domain.

*optn* ΔLIR cDNA was amplified from *optn*-pCS2+ constructs by designed primers (S5 Table), excluding the LIR protein domain. The PCR products were gel purified by Quick gel Extraction PCR Purification Combo Kit (K220001,Invitrogen) and the two fragments and pCS2+ plasmid were digested by BamH1(R0136S,NEB) and EcoR1(R0101S,NEB), after which the two fragments were ligated into pCS2+ plasmid by T4 DNA ligase.

*p62* ΔUBA cDNA was obtained from a *p62*-pCS2+ construct by Nco1(R0193S, NEB) digestion and religation, which excludes the region encoding the UBA protein domain.

*p62* ΔLIR cDNA was obtained from a *p62*-pCS2+ construct by NcoN1 digestion and religation.

*Optn* mRNA,*optn* ΔUBAN, and *optn* ΔLIR mRNA was generated using SP6 mMessage mMachine kit (Life Technologies) from Kpn1 or Sac1(R0156S, NEB) digested *optn*–pCS2+ constructs. RNA purification was performed using the RNeasy Mini Elute Clean up kit (QIAGEN Benelux B.V., Venlo, Netherlands).

*In vitro* transcription of *p62*, *p62* ΔUBA, and *p62* ΔLIR was performed using mMESSAGE mMACHINE® T3 Transcription Kit (AM1348, Thermo Fisher) and purified using the RNeasy MiniElute Cleanup kit (QIAGEN Benelux B.V., Venlo, Netherlands). All mRNAs were injected into one cell stage embryos, and the overexpression effects of *optn* or *p62* were validated by Q-PCR and Western blot.

### Gene Expression Analysis

Total RNA was extracted using Trizol reagent (15596026, Invitrogen) according to the manufacturer’s instructions and purified with RNeasy Min Elute Clean up kit (Lot:154015861, QIAGEN). RNAs were quantified using a NanoDrop 2000c instrument (Thermo Scientific, U.S). Reverse transcription reaction was performed using 0.5 µg of total RNA with iScript cDNA synthesis kit (Cat:#170-8891, Bio-Rad). The mRNA expression level was determined by quantitative real-time PCR using iQSYBR Green Supermix (Cat:170-8882, Rio-Rad) and Single color Real-Time PCR Detection System (Bio-Rad, U.S) as previously described (50). All primers are listed in S5 Table.

### Statistical analyses

Statistical analyses were performed using GraphPad Prism software (Version 5.01; GraphPad). All experimental data (mean ± SEM) was analyzed using unpaired, two-tailed t-tests for comparisons between two groups and one-way ANOVA with Tukey’s multiple comparison methods as a posthoc test for comparisons between more than two groups. (ns, no significant difference; *p < 0.05; **p < 0.01; ***p < 0.001). To determine whether the offspring of F1 heterozygous mutants follows Mendelian segregation, the obtained data was analysed with a Chi-square test (ns, no significant difference).

## Acknowledgements

We thank Daniel Klionsky for sharing of the GFP-Lc3 transgenic zebrafish line. We are grateful to all members of the fish facility team for zebrafish caretaking. We would like to thank Gerda Lamers and Joost Willemse for advice on confocal imaging and image analysis.

## Supplementary Figure Legends

**S1 Fig. Optineurin and p62 are highly conserved between zebrafish and human**

(A) Representative images of *WT* and mutant F2 embryos at 4dpf. Scale bars, 250 µm.

(B) Phylogenetic tree of SLR amino acid sequences. Optineurin, p62, NDP52(Calcoco2), NBRC1 and TAX1BP1 sequences were searched from the NCBI Ensembl database and the accession numbers are listed in S6 Table. MUSCLE online server was used to generate the protein alignment. The best-fitting amino acid replacement model to the alignment (JTT) was determined using ProtTest 3.2 based on the Akaike Information Criterion (AIC). Finally, the maximum likelihood gene tree was estimated with PhyML 3.0 and represented in FigTree v1.3.1 (http://tree.bio.ed.ac.uk/software/figtree/). Nodal confidence was calculated with non-parametric bootstrap of 100 replicates.

(C) Protein sequence identity of SLRs between zebrafish and human. The percentage identity and similarity was calculated using a Clustal Omega alignment.

(D) Alignment of LIR, UBAN and UBA motifs from the Optn and p62 sequences of different vertebrates. Amino acid sequences of the LIR motifs of Optn and p62 from the indicated species were aligned using Mega7 software (DNASTAR, Madison, WI) and aligned by the Clustal W2 method (EMBL, Cambridge, UK). The Ubiquitin binding domains of Optineurin or p62 were determined by NCBI-BlASTP (https://blast.ncbi.nlm.nih.gov/Blast.cgi?PAGE=ProTeins).

**S2 Fig. Characterization of Optineurin and p62 mutant lines**

(A) Validation of Baf A1 effect on zebrafish by Western blot. Baf A1 treatment at dosages of 20, 100 and 400 nM was performed by incubation for 12h in egg water. The protein samples were extracted from 4 dpf AB/TL larvae (>10 embryos/sample). The blots were probed with antibodies against Lc3 and Actin. (B) Detection of p62 or Optineurin protein in mutant lines in absence or presence of Baf A1. Protein samples were extracted from *optn*^+/+^, *optn*^Δ5n/Δ5n^, *p62*^+/+^ and *p62*^Δ37n/Δ37n^ larvae at 4 dpf (>10 embryos/sample). The blots were probed with antibodies against Optineurin, p62 and Actin as a loading control. Optineurin/Actin and p62/Actin ratios are indicated below.

**S3 Fig. Injection of *optn* or *p62* MO transiently knocks down the corresponding mRNA and protein.**

(A) Workflow representing the experimental design in (B-E). *optn* or *p62* MO were injected into one cell stage embryos (AB/TL), and injected embryos were collected for confirmation of the knockdown effect by RT-PCR and Western blot analysis (>20 embryos /Sample).

(B) Validation of the effect of *optn* splice-blocking MO e2i2 (targeting the splice event between exon 2 and intron 2) by RT-PCR on (a) the wild type control group, (b) embryos injected with 0.1mM MO, or (c) embryos injected with 0.15 mM MO. The wild type PCR product is expected to be 400 bp in length.

(C) Validation of the effect of *p62* splice-blocking MO i1 e2 (targeting the splice event between intron 1 and exon 2) by RT-PCR on (a) the wild type control group, (b) embryos injected with 0.5mM MO. The wild type PCR product is expected to be 200 bp in length.

(D and E) Validation of MO knockdown effect by Western blot analysis. The protein samples were exacted from 2 dpf AB/TL embryos injected with 0.1mM *optn* or 0.5 mM *p62* MO (>20 embryos/sample). The blots were probed with antibodies against Optn or P62 and Actin.

**S4 Fig. Optineurin or p62 mutation reduces autophagosome formation during Mm infection**

(A) Representative confocal micrographs of GFP-Lc3 co-localization with Mm clusters in *optn*^+/+^, *optn*^Δ5n/Δ5n^, *p62*^+/+^ and *p62*^Δ37n/Δ37n^ infected embryos at 1 dpi. The arrowheads indicate the overlap between GFP-Lc3 and Mm clusters. Scale bars, 10 μm.

(B) Quantification of the percentage of Mm co-localizing with GFP-Lc3 in infected embryos at 1dpi (>6 embryo/group). ns, non-significant, *p<0.05,**P<0.01,***p<0.001. (C) Autophagy activity in Mm infected embryos. Protein samples were obtained from 3 dpi *optn*^+/+^, *optn*^Δ5n/Δ5n^, *p62*^+/+^ and *p62*^Δ37n/Δ37n^ infected larvae with Baf A1 12 h treatment (>10 larvae/sample). The blots were probed with antibodies against Lc3 and Actin.

**S5 Fig. Transient overexpression of *optn* or *p62* mRNA reduces the susceptibility to Mm**

(A,B) Quantification of Mm infection burden at 2dpi in embryos injected with full length or ΔLIR/ΔUBAN deletion mRNAs of *optn* (A) and *p62* (B). Data are accumulated data from two independent infection experiments. ns, non-significant, *p<0.05,**P<0.01,***p<0.001.

**S6 Fig. Transient overexpression of *optn* or *p62* mRNA results in increased recruitment of GFP-Lc3 to Mm clusters at 1 dpi.**

(A) Representative confocal micrographs of GFP-Lc3 co-localization with Mm clusters in mRNA-injected larvae at 1 dpi. The arrowheads indicate the overlap between GFP-Lc3 and Mm clusters. Scale bars, 10 μm.

(B) Quantification of the percentage of Mm clusters positive for GFP-Lc3 vesicles. ns, non-significant,*p<0.05,**P<0.01, *** p<0.001. Data are accumulated from two independent experiments (>15embryo/group).

Supplementary Tables

S1 Table. Zebrafish lines used

S2 Table. Target sites for CRISPR/Cas 9 systems

S3 Table. Primers for complementation and amplification of sgRNA

S4 Table. Morpholino sequences

S5 Table. Primers used in this study

S6 Table. Accession numbers of selective autophagy receptors

